# Ribosomal dysregulation: A conserved pathophysiological mechanism in human depression and mouse chronic stress

**DOI:** 10.1101/2023.05.04.539419

**Authors:** Xiaolu Zhang, Mahmoud Ali Eladawi, William George Ryan, Xiaoming Fan, Stephen Prevoznik, Trupti Devale, Barkha Ramnani, Krishnamurthy Malathi, Etienne Sibille, Robert Mccullumsmith, Toshifumi Tomoda, Rammohan Shukla

## Abstract

The underlying biological mechanisms that contribute to the heterogeneity of major depressive disorder (MDD) presentation remain poorly understood, highlighting the need for a conceptual framework that can explain this variability and bridge the gap between animal models and clinical endpoints. Here, we hypothesize that comparative analysis of molecular data from different experimental systems of chronic stress and MDD has the potential to provide insight into these mechanisms and address this gap. Thus, we compared transcriptomic profiles of brain tissue from postmortem MDD subjects and from mice exposed to chronic variable stress (CVS) to identify orthologous genes. Ribosomal protein genes (RPGs) were downregulated, and associated RP-pseudogenes were upregulated in both conditions. Analysis across independent cohorts confirmed that this dysregulation was specific to the prefrontal cortex of both species. A seeded gene co-expression analysis using altered RPGs common between the MDD and CVS groups revealed that downregulated RPGs homeostatically regulated the synaptic changes in both groups through a RP-pseudogene-driven mechanism. *In-vitro* and *in-silico* analysis further demonstrated that the inverse RPG/RP-pseudogene association was a glucocorticoid-driven endocrine response to stress that was reversed during remission from MDD. This study provides the first evidence that ribosomal dysregulation during stress is a conserved phenotype in human MDD and CVS exposed mouse. Our results establish a foundation for the hypothesis that stress-induced alterations in RPGs and, consequently, ribosomes contribute to the synaptic dysregulation underlying MDD and chronic stress-related mood disorders. We discuss a ribosome-dependent mechanism for the variable presentations of depression and other mood disorders.

**Significance Statement:** The presented study highlights the pressing need for a connection between animal models of depression and clinical endpoints. The lack of concordance between these two areas has hindered our understanding of MDD’s biological underpinnings. The study’s hypothesis that orthologous gene from experimental systems of chronic stress and MDD can bridge this gap is a major advance in this field. The study indicates that dysregulation of ribosomes in the synapse is a common feature in both human MDD and mice exposed to CVS. This dysregulation is a response to endocrine disturbances and is driven by mechanisms that involve pseudogenes. These findings support the hypothesis that stress-induced alterations in RPGs and, consequently, ribosomes may contribute to the variable presentations of depression.

## Introduction

Major depressive disorder (MDD) is a debilitating mental illness that affects more than 300 million people worldwide and is ∼2 times more prevalent in females. Therapies and medication are available, but a 13% increased prevalence of MDD in the past decade suggests limited success of these treatments. This shortcoming between therapies and remission can be linked to the heterogenous nature of depression presentation and our inadequate understanding of the disease biology. Human postmortem brain research has provided important insight in disease biology but is often associated with uncontrollable biological and environmental variables and undocumented medical histories. Thus, animal models of depression are a crucial tool for examining several molecular and cellular changes associated with depression in a controlled environment (1). For example, in rodents, chronic variable stress (CVS), a paradigm involving systematic and repeated exposures to variable unpredictable and uncontrolled stressors over days or weeks, recapitulates several features of MDD (2). However, in spite of the advances this (and other (1)) models have facilitated, a translational gap remains, largely due to the poor correlation between the aspect of depression explained by animal models and clinical endpoints. Many traits of human depression are either absent, difficult to assess, or radically different in animal models. Prior transcriptomics-based studies have attempted to identify similarities between rodent CVS and human MDD using integrated network-based analysis to identify a common hub gene for downstream analysis (3). However, such strategies often fail to reproduce reliable associations with disease (4). We hypothesize that identifying ortholog gene families, rather than hub genes, that are similarly affected between human MDD, and mouse CVS will provide seeds for gene network analyses generating ontologies that uncover structural and biochemical foundations of disease pathophysiology. Importantly, this strategy can be replicated using other datasets to highlight similarities and differences between species or sexes to further our understanding of the biological basis of disease heterogeneity and sexual dimorphism.

We initiated this study by identifying orthologous gene families altered in both the human MDD and mouse CVS condition in existing (3) total RNAseq data from the prefrontal cortex (PFC; dorsolateral prefrontal cortex (DLPFC) in human). Ribosomal protein genes (RPGs)—an evolutionary conserved family of genes—showed significant overlap between the two species in a region- and cell-compartment-specific manner. Our findings provide a foundation for a hypothesis proposing that stress-induced alterations in RPGs, which occur in species ranging from yeast to rodents and humans, result in ribosome dysregulation that may underlie pathophysiological changes observed in chronic stress and stress-related mood disorders. Importantly, the diverse infrastructure and mobility of ribosomes across neurites provide an opportunity for ribosome dysregulation to explain the immense variability in stress-response phenotype and human depression subtypes.

## Results

### RPGs are dysregulated in the PFC of CVS-exposed mice and humans with MDD

To identify gene family orthologs that are conserved in the DLPFC of humans with MDD and PFC of mice exposed to 4 weeks CVS, we performed differential expression analysis (Table. S1) on reposited transcriptome datasets from previous work (3) and examined the enrichment of ∼1200 HGNC (Human Genome Nomenclature Committee (5)) gene families in the sets of up- and downregulated differentially expressed genes. We discovered significant enrichment of gene families associated with large and small RPGs in the downregulated genes of male and female mice and male humans (Fig. 1A, Table. S2). After harmonizing the gene symbols between the two species, we examined the overlap in the significant downregulated genes and identified 15 RPGs that were differentially expressed in the DLPFC of humans with MDD and PFC of mice exposed to CVS (Fig. 1B, common RPGs henceforth).

**Fig. 1:**
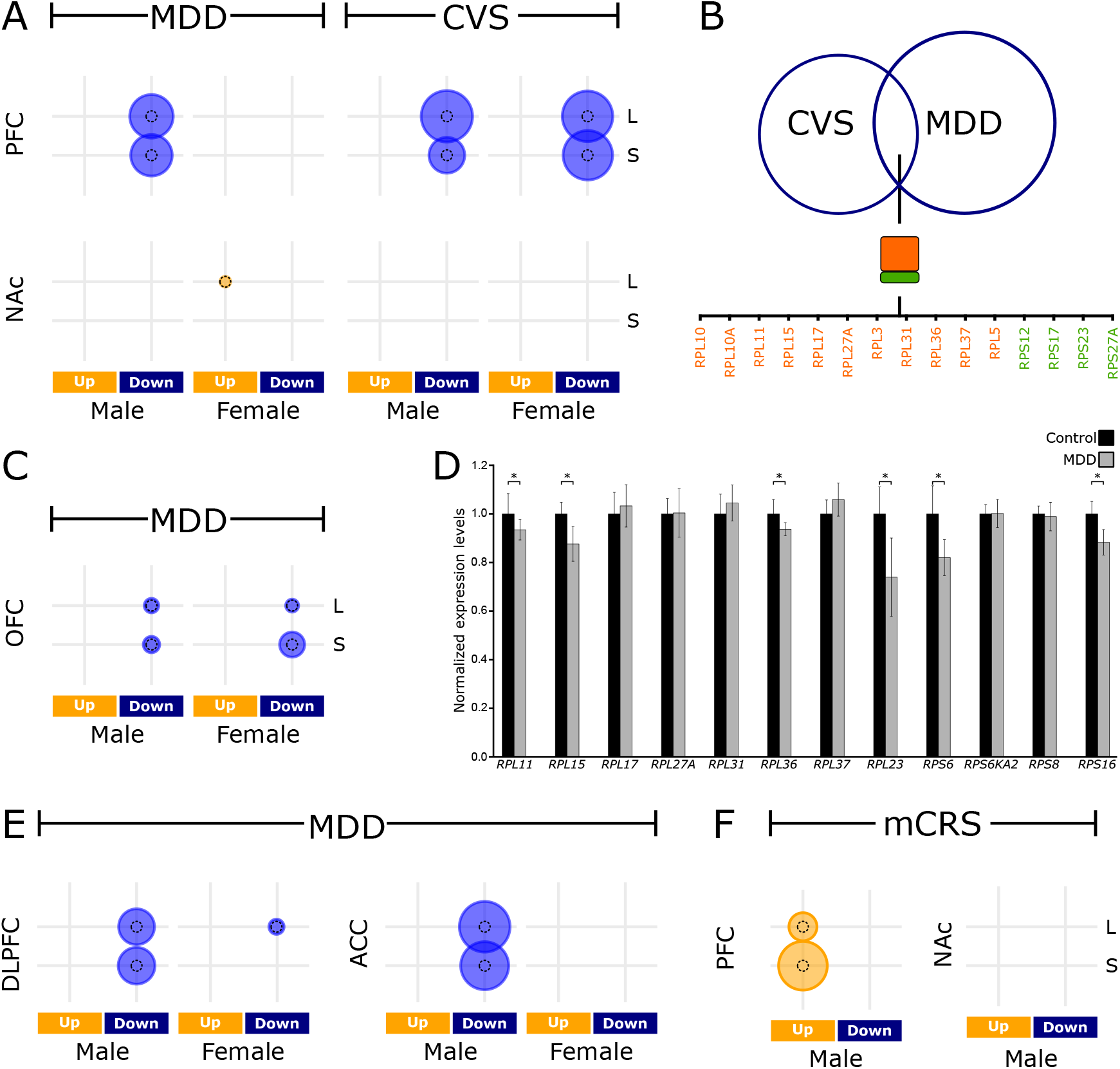
RPGs in the PFC are dysregulated during MDD and CVS, **A)** Enrichment of large (L) and small (S) subunit RPGs in the downregulated genes in the PFC and NAc of humans with MDD and CVS-exposed mice. **B)** Overlap of downregulated RPGs in MDD and CVS. **C)** Enrichment of RPGs in the OFC from same cohort. **D)** qPCR results showing 6/12 common RPGs are downregulated in the ACC of an independent cohort of MDD patients. n = 6/group; * < 0.05. **E&F)** RPGs were dysregulated during MDD **(E)** and mCRS **(F)** in prefronatal cortex data avilable from independent studies. Size of the circle is proportionl to -log_10_(q-value). The reference dotted circle is proprtional to -log_10_(q-value = 0.05).

Prior studies have shown that RPG expression varies by tissue type (6). Thus, we investigated the enrichment of large and small RPG families in the transcriptomic data for the nucleus acumens (NAc, Table. S1) of both species from the same cohort and analysed additional data available from the frontal cortex region comprising the orbitofrontal cortex (OFC, Table. S1) in the same human cohort. The NAc of neither species showed enrichment of RPG families (Fig. 1A, bottom; Table. S2). However, we did observe a significant enrichment of the RPG family in the downregulated genes of the OFC for both males and females, albeit a less significant enrichment than the one observed in the DLPFC (Fig. 1C, Table. S2). qPCR analysis of common RPGs in the anterior cingulate cortex (ACC) of male MDD and control subjects from an independent cohort also revealed significant downregulation of several common RPGs in the ACC (Fig. 1D). Similarly, analysis of independent reposited datasets (7) from the DLPFC and ACC of humans with MDD confirmed a significant enrichment of RPGs in downregulated genes in both regions (Fig. 1E, Table. S1 & S2).

To further validate our results in mice, we examined the differential expression profile of an independent dataset from the PFC and NAc of mice who had undergone 3 weeks of multimodal chronic restrain stress (8) (mCRS; Fig. 1F), a stress paradigm with a different stress regime and allostatic load than the CVS paradigm. Consistent with the CVS results, enrichment analysis showed that RPGs were significantly dysregulated in the PFC but not in the NAc in mCRS-exposed mice (Fig. 1F, Table. S2). However, unlike the CVS-exposed mice, the mCRS-exposed mice had upregulated RPGs.

Overall, evidence from multiple independent studies demonstrates that gene families associated with ribosome structure are variably (i.e., either up or down) but significantly and consistently dysregulated during stress in mice and MDD in humans, with the dysregulation primarily occurring in frontal cortical regions in both species. As ribosomal proteins (RPs) constitute the ribosomes, these results point towards altered translation and translational machinery during stress.

### Seeded gene co-expression analysis reveals species- and sex-specific ribosome regulation

Next, we reasoned that, in a mechanism analogous to hub genes in gene co-expression network analysis, we could treat the common RPGs as seeds to identify correlated genes (spearman correlation, p-value < 0.05; Table. S3), thereby identifying the biological processes coordinated by the RPGs. Furthermore, we hypothesized that stratifying this seeded gene co-expression analysis for all conditions (i.e., stress/disease and control state in both sexes of both species) would reveal important sex and species-specific differences in RPG-dependent regulation. The Gene Ontology (GO)-terms associated with the RPG-correlated genes (q-value < 0.05) across all conditions were clustered into different themes (Fig. 2, left labels; Table. S3), established in our previous meta-analysis of different psychiatric disorders (9, 10). Several intriguing associations stood out:

**Fig 2:**
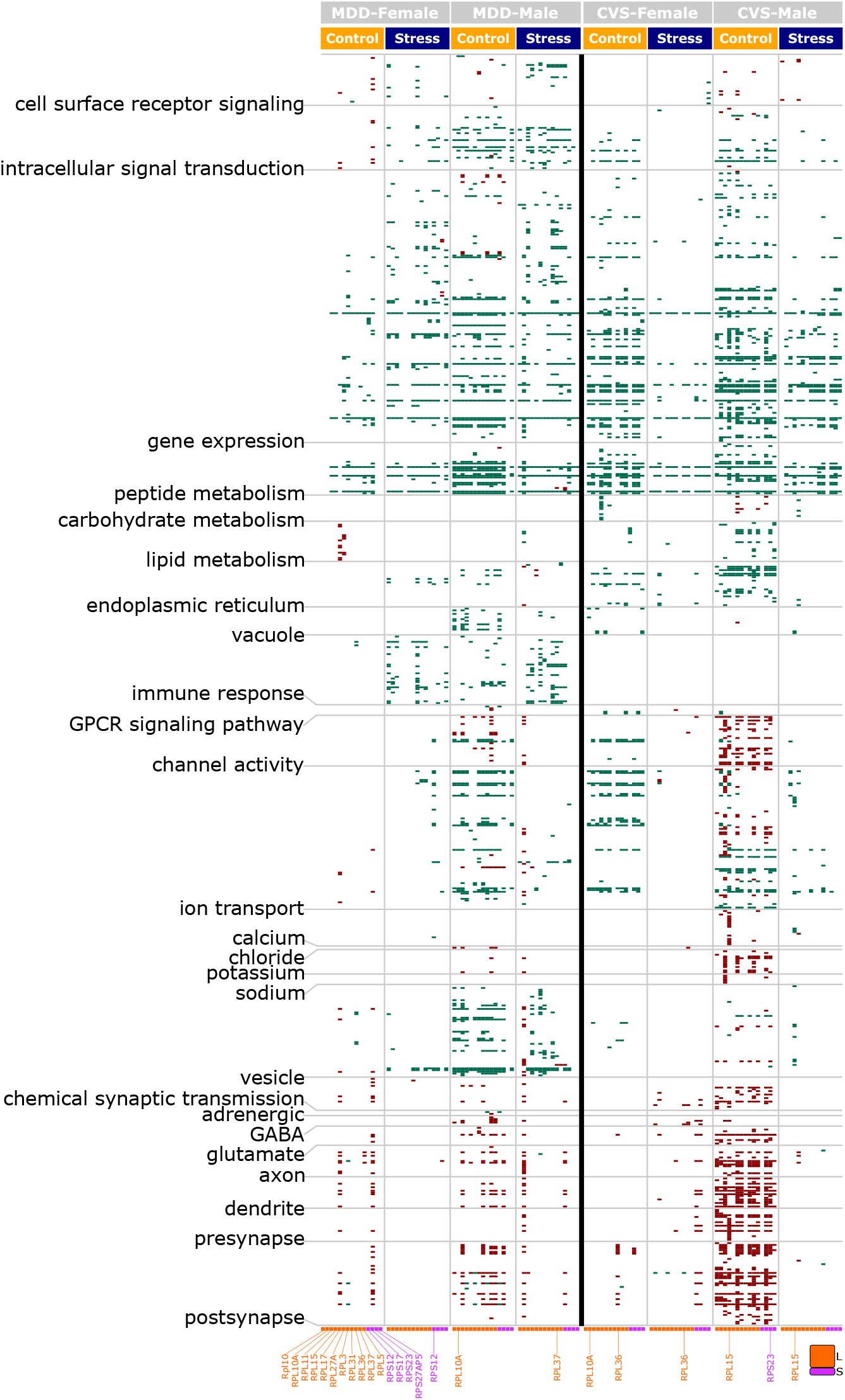
The MDD- and CVS-sentive changes in RPGs inversely regulate pathways associated with synaptic functions. A heatmap of common RPG-regulated pathways (q < 0.05) in both sexes during MDD and CVS phenotypes under control and stress conditions. Green and maroon: pathways positively and negatively regulated by RPGs, respectively. Notice the negative correlation of RPGs with intercellular communication related themes and the variable number of RPGs associated with the control and stress state.

First, RPG-based regulation showed a striking contrast in the themes associated with positive and negative (shown as green and maroon, respectively) correlation. Nearly all intracellular events involving signal response coupling (*cell surface receptor signaling, intracellular signal transduction, and gene expression*), metabolic processes (*metabolism of peptides, carbohydrates, and lipids*), cellular organelles (*endoplasmic reticulum and vacuole*), and the immune response were positively correlated with RPGs. On the other hand, intercellular signaling events involving neuromodulatory infrastructure (*calcium, chloride, potassium, sodium, adrenergic, GABA and glutamate*), neuromodulation activity (*G protein-coupled receptor [GPCR] signaling, channel activity, ion transport, chemical synaptic transmission*), and synaptic infrastructure (*vesicles, axon, dendrite, presynapse and post synapse*) were negatively correlated with RPGs. Notably, a negative correlation reflects a system’s homeostatic balance (11), with inhibition of some genes and pathways being associated with the stimulation of others. Thus, the negative correlation of RPGs with synaptic alterations suggests a role for these common RPGs in homeostatic modulation of synaptic input and output.

Second, themes and pathways identified in the control state but absent in the MDD or CVS states suggest RPG coordination that was lost with stress or disease, and themes and pathways absent in the control state but present in the MDD or CVS states suggest new stress-altered coordination. Accordingly, females with MDD demonstrated increased intracellular events and decreased intercellular events related to RPG coordination compared to males, who exhibited only a marginal decrease in coordination across all themes in the MDD state. In CVS-exposed mice however, this pattern differed and both female and male mice exhibited decreased coordination across all themes in the stressed state. Notably, there was increased RPG coordination of the immune response in both sexes in the MDD population than in the controls, but no significant coordination of the immune response with RPGs was observed in CVS-exposed mice.

Lastly, among 15 common RPGs (Fig. 1B) belonging to either the large or small subunit used for the analysis, a variable number and type (large or small) of RPGs were associated with the control and stress/disease states. Interestingly, the number of RPGs contributing to negatively correlated themes involving neuromodulatory infrastructure (*calcium, potassium, sodium, GABA, and glutamate*), fast neuromodulation activity (*channel activity, ion-transport, chemical synaptic transmission*) and synaptic infrastructure (*vesicles, axon, dendrite, presynapse and post synapse*) were decreased in the stress/disease condition in sex- and species-specific manners. For example, there was a decreased coordination of these themes in CVS-exposed male mice than in control males.

Overall, the findings using the seeded gene network analysis imply that RPGs are responsible for the homeostatic feedback regulation of pathways associated with synaptic communication during both stress and MDD in both sexes. However, this regulation is differentially influenced by a diverse set of RPGs, indicating that ribosome composition may be changed in a sex- and species-specific manner.

### RPG dysregulation is likely governed by RP-pseudogenes

RP-pseudogenes are evolutionarily mutated DNA sequences that resemble RPGs and often regulate the expression and function of the parent RPGs. At the transcript level, regulation is either through RNA interference, where a pseudogene serves as a short interfering RNA (siRNA) to downregulate parent RPG expression, or through the competitive endogenous RNA (ceRNA) pathway, where a pseudogene acts as a sponge for common micro RNAs (miRNAs) to attenuate downregulation of parent RPG expression (12, 13). Notably, in mice and humans, the majority of downregulated RPG parent genes had upregulated expression of RP-pseudogenes (Fig. 3, top), implying that the majority of the observed downregulated RPGs are regulated via their pseudogenes. To determine the RPG-associated functions dependent or independent of RP-pseudogenes, we mapped the ontologies (q-value < 0.05) associated with downregulated RPGs (Fig. 3, maroon columns) and upregulated RP-pseudogenes (Fig. 3, green columns) to each other. There were phenotype-specific (CVS or MDD) differences in themes (Fig. 3, middle labels) associated with autophagy, extracellular region (ECR), signaling, organelle, and protein metabolism, and these themes showed minimal overlap with RPG- and RP-pseudogene-associated functions. However, canonical functions associated with ribosomes and RNA, as well as most synaptic pathways, revealed considerable overlap in RPG- and RP-pseudogene-associated functions during MDD and CVS, indicating that RP-pseudogenes can selectively modify these RP-related functions in both species during stress and MDD.

**Fig. 3:**
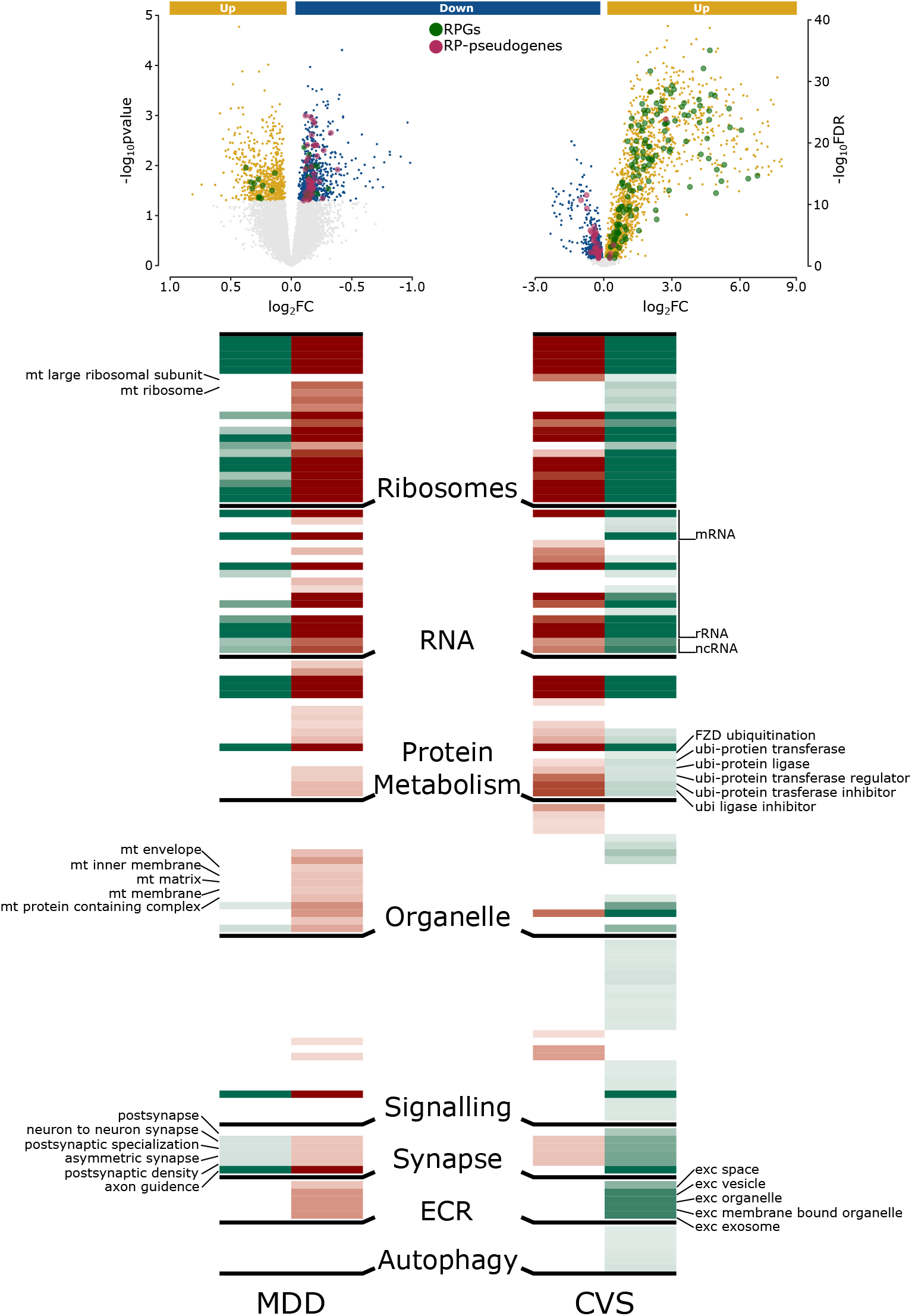
RPG downregulation is regulated by pseudogenes: Pathway profile associated with the downregulated RPGs (maroon) and upregulated RP-pseudogenes (green) is shown. The pathways associated with both the RPG and RP-pseudogene are likely to be linked with an RP-pseudogenes during the stressed state. Notice that most of the housekeeping-(*Ribosomes, RNA, and Protein metabolism*) and synapse-associated pathways associated with RPGs overlap with RP-pseudogenes. The lighter to darker shade of both maroon and green show increasing enrichment calculated as -log_10_(*q-value*).

Overall, RPG and RP-pseudogene-specific functional analysis suggests that several of the RPG functions are regulated by their respective pseudogenes. Additionally, the pseudogene regulation of intracellular singling events involving the synapse may explain the observed negative correlation of these event with RPG expression (Fig. 2).

### Remission from MDD is associated with RPG downregulation reversal

Next, we reasoned that if RPG dysregulation is functionally associated with MDD, RPG downregulation should be reversed during remission from MDD. To probe this hypothesis, we examined our existing datasets (14) that included data from the ACC of postmortem subjects who died during MDD episodes or in remission. Consistent with our core results, we observed downregulated RPGs and upregulated RP-pseudogenes during MDD episodes. However, during remission, the downregulated RPG and upregulated RP-pseudogene relationships observed during an MDD episode were reversed (Fig. S1). Furthermore, the canonical ribosome-related pathways (q-value < 0.05) were downregulated during episodes and upregulated during remission. To validate the accuracy of the episode and remission state, we examined their characteristic alterations. Episode state, consistent across several previous reports, was associated with downregulated pre- and post-synaptic changes (15) and upregulated glucocorticoid and immune response (16). Likewise, the episode and remission states, which are both defined as a depression trait, showed upregulation of the innate immune response (14).

Overall, independent analysis of additional datasets confirms the inverse association between RPG and RP-pseudogenes, and analysis of the remission state indicates its functional association with MDD.

### RPGs are dysregulated in glucocorticoid-treated primary PFC neurons

Both MDD (Fig. S1) and CVS (17) are associated with elevated levels of glucocorticoid activity, which are responsible for several of the negative consequences of chronic stress. Therefore, we reasoned that chronic treatment (72 h) (18, 19) of primary PFC neurons with a synthetic glucocorticoid, dexamethasone (DEX, 1 μM) (18), may replicate the inverse dysregulation of RPG and RP-pseudogene expression observed during MDD and CVS. To verify stress induction in our *in vitro* model, we established that DEX-treated PFC cells showed prominent formation of stress granules (SGs), a putative marker of stress response (Fig. 4A & B) (20, 21). Additionally, qPCR analysis demonstrated that DEX-treated PFC cells expressed several known markers of chronic stress identified in experimental models (Fig. 4C-G) (22-27). We then compared the RNAseq-based expression profiles of control and DEX-treated primary neurons. Notably, consistent with the expression profiles of the MDD patients and CVS-exposed mice (Fig. 3 top and Fig. S1), we observed an inverse association between RPGs and RP-pseudogenes in DEX-treated primary neurons that was attenuated when the cells were co-treated with DEX and RU_486_, a glucocorticoid receptor antagonist (Fig. 4H). Importantly, the upregulated RP-pseudogenes influenced pathways related to synaptic signalling themes (Fig. 4I).

**Fig. 4:**
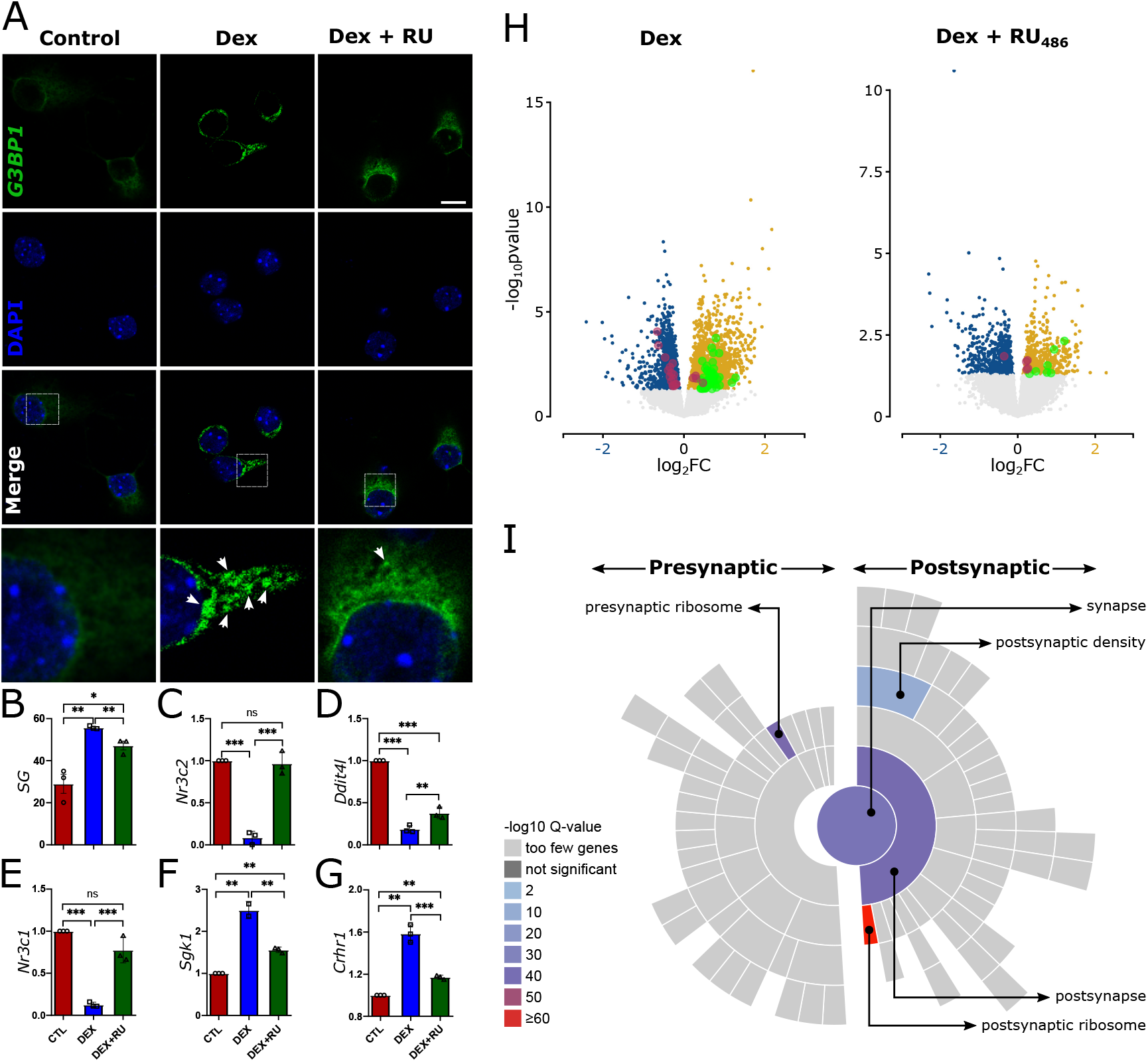
The inverse association between RPG and RPG pseudogenes is preserved in PFC primary neurons. **A)** DEX-treated PFC primary neurons displayed SG expression (white arrow), which is suppressed by RU_486_ treatment. **B)** Quantification of SG expression in control DEX and DEX + RU_486_ cells. Scale bar: 25 μm. **C-to-G)** The results of qPCR demonstrate that CVS markers are similarly dysregulated in glucocorticoid-stressed neuronal cells. n = 3/group **H)** The inverse association between RPG and RPG pseudogenes observed during DEX is reversed is lost after RU_486_ treatment. n = 12/group **I)** Similar to RP-pseudogenes in the CVS and MDD conditions, the RP-pseudogenes were significantly associated with synaptic pathways.

Overall, the *in vitro* experiment confirms the chronic stress-related endocrine origin of RPG dysregulation. Thus, across all three chronic stress paradigms (human MDD, mouse CVS and glucocorticoid stressed primary neurons), pseudogenes may play a vital role in regulating stress response pathways and synapse-associated RPG functionality.

## Discussion

The RPG family, with > 110 members, is a highly diverse group of genes that serve as the building blocks of ribosomes. Intriguingly, ribosomes are involved in the stress response across several phyla, with the stress-related roles of ribosomes being most studied in simple life forms such as bacteria and yeast. Here, in accordance with their evolutionarily conserved involvement with stress, we demonstrated involvement of RPGs in human MDD and mouse chronic stress and confirmed our results using qPCR analysis of multiple datasets and independent cohorts. Consistent with the known region-specific distribution of RPs (6), stress/depression-related RPG dysregulation was observed in the PFC of both humans and mice but not in the NAc. The neurons of the PFC have a high density of synapses,(28) which require large amounts of ribosome machinery. Thus, the enrichment of differentially expressed RPGs is not surprising. However, the seeded gene-correlation analysis (Fig. 2) revealed that the differentially expressed RPGs are positively correlated with soma-related biological processes and negatively correlated with neurite-related processes. The negative correlation is suggestive of a homeostatic feedback loop (11). Accordingly, we provide evidence that the negative correlation of RPGs with neurite-related processes may be governed by RP-pseudogenes, which can regulate RPG expression at the transcript level. Finally, we observed that the number and type of RPGs differentially expressed in the stress/disease and control states varied by sex. Ribosomes are heterogenous, with numerous possible permutations in RP composition. Based on these findings, we posit a ribosome hypothesis of depression: Ribosomes, via their pseudogene-regulated diverse infrastructure and mobility across neurites, may provide regulatory diversity that tunes information input and output. Our evidence suggests that both human MDD and mouse chronic stress are associated with a dysregulation of RPGs and ribosome composition in neurites of the PFC, potentially providing a pathophysiological basis for the synaptic dysregulation underlying MDD and the chronic stress response. Considering our functional analysis, we will discuss how differential regulation of RPGs may be related to stress and stress-related disorders such as MDD.

To encode and learn complex patterns, each cortical neuron must access many possible combinations of axonal input (Fig. S2). These inputs can target various parts of the neuron, including the soma, distal dendrites, and proximal dendrites. Several psychiatric disorders are linked to location-specific dysregulation of inputs. For instance, dendritic input is reportedly altered during MDD (10), and somatodendritic input during schizophrenia (29). Our hypothesis posits that ribosomes, along with their diverse RPs, sculpt the adaptability of neurons to variable inputs through the following potential mechanisms (Fig. S2). First, location-non-specific downregulation of RPGs can lead to decreased ribosome biosynthesis, thus reducing global translation and protein synthesis. Ribosome biosynthesis is an energy-intensive process that requires several essential amino acids and the presence of adenosine triphosphate (ATP). Consequently, even simple species such as *saccharomyces cerevisiae* have a homeostatic mechanism during stress that halts ribosome biosynthesis via downregulation of RPGs to conserve energy (30). RPG downregulation in human MDD and mouse CVS may serve to preserve several essential amino acids, such as lysine and arginine, which are abundant in ribosomes (31, 32) and whose deficiency has been functionally linked to depression (33-35). In neurons, most RPGs are found in neurites; thus, the homeostatic downregulation of RPG will decrease synaptic protein synthesis. Further support for the role of RPG downregulation in maintaining synaptic homeostasis comes from the negative coordination of common RPGs with most intercellular singling-related themes, which was decreased in the stress and disease conditions (Fig. 2). Interestingly, this appears to be consistent with the process of synaptic scaling, a plasticity mechanism in which a neuron regulates its own excitability in relation to network activity, primarily by targeting the protein synthesis process. Indeed, males in the human MDD and mouse CVS groups exhibited an excessive loss in coordination of pathways associated with glutamatergic singling (Fig. 2). The mechanisms of homeostatic synaptic scaling are largely unknown. Our results point toward RP-pseudogenes as potential candidates driving the homeostatic downregulation of RPGs (Fig. 3). Notably, unlike long-term potentiation or depression, where synaptic strengths are rapidly potentiated or depressed, synaptic scaling develops over several hours to days (36) and is dependent on protein synthesis. The average half-life of pseudogenes (10 to 17h) (37) falls into the range of duration required for homeostatic scaling, further supporting the possible role of RP-pseudogenes in mechanisms of homeostatic synaptic scaling.

Homeostatic plasticity can operate at a local level, by targeting a group of synapses on a certain dendritic branch, or at a synapse-specific level (38). Following that, the second mechanism by which ribosomes can sculpt the adaptability of neurons to variable inputs is through a site-specific modification of ribosome composition. Ribosome heterogeneity allows for specialized ribosomes that provide preferential translation regulation (39). Specialized ribosomes can have distinct modifications with functional outcomes ranging from changes in translation initiation, speed, fidelity control, and mRNA translation selectivity (Fig. S2). Functional specializations of ribosomes may also result from activity-dependent phosphorylation (40) and other post-translation modification (41) on selected RPs, thereby further modifying ribosome composition. Future studies will explore stress-induced ribosome specialization and the functional consequences on neuronal input and output and synaptic dysregulation.

These two mechanisms by which ribosomes can sculpt the adaptability of neurons to variable inputs—location-non-specific RPG downregulation and site-specific modification of ribosome composition—have several implications for mood disorders and therapeutic strategies targeting them. First, depression can be viewed as part of a spectrum of mood disorders, with several types of depression lying along a continuum with nebulous boundaries between them. Dendritic input is dysregulated during depression, resulting in a remodelling of the microcircuitry associated with learning, memory, and attention (10). Together, location-non-specific RPG downregulation and site-specific modification of ribosome composition can result in several permutations supporting dysregulation of synaptic input along the dendritic arbor. These permutations can contribute to nuances resulting in the spectrum nature of depression and other mood disorders. Second, targeting RP expression regulation and its role in ribosomal heterogeneity offers a promising therapeutic strategy previously unexplored. Ribosomes are typically associated with protein synthesis, which in dendrites can involve transmitter receptors, synaptic scaffold proteins, and other regulatory proteins. Thus, our novel ribosome hypothesis can encompass other known hypotheses of depression involving GABA (42), glutamate (43), immune (44), and monoaminergic (45) systems. As such, exploring the therapeutic options associated with ribosomal dysregulation will have a potentially broader transdiagnostic impact across depression subtypes.

## Materials and Methods

### Data cohorts

We downloaded datasets in the fastq.gz format from Gene Expression Omnibus (GEO) with ID GSE102556 (3), which included both mouse and human datasets for CVS and MDD, respectively. Mouse data was available only from PFC (female=19, male=19) and NAc (female=20, male=20). For humans, data was available from several brain regions, but we focused on analysing the regions homologous to the available mouse regions. This included DLPFC (female=22, male=26, mean age = 47) and OFC (female=22, male=26, mean age = 47) and basal for brain region involving NAc (female=22, male=28, mean age = 47). To validate our findings, we used additional datasets from mouse subjected to multimodal chronic restrain stress (mCRS) paradigm (GSE148629) and human MDD (GSE80655)^8^. GSE148629 had both mouse PFC (n=15, age = 15 weeks) and NAc (n=8, age = 15 weeks) data from male. GSE806558 had MDD data from human DLPFC (female=6, male=17, average age = 46 years) and ACC (female=6, male=16, average age = 45 years), which is also a prefrontal cortex region. We validated the ACC related findings using our previous study (14) and quantitative polymerase chain reaction (qPCR) analysis described in the subsequent methods section.

### Alignment Protocol

We used HISAT2 version 2.1.0 to index mouse and human reference genomes and align downloaded RNAseq data. We then used subread version 1.5.0-p2 to quantify reads and map to genomic features. Notably, RPGs have about 2000 RP-pseudogenes that are significantly homologous to the parent RPGs (46), which impedes the ability of the standard alignment method to uniquely identify reads mapped to RPG and RP-pseudogenes with high sequence homology. To circumvent this, we used a specialized alignment method as described before (47). Briefly, we mapped all sequence reads to a “composite genome” that includes the entire human (GRCh38) or mouse (GRCm38) genome sequence and spliced mRNA sequences of RPGs (47). We then mapped RNA-seq reads to the composite genome without allowing mismatches and discarding reads mapped to more than one locus, thus ensuring that the reads mapped to RP-pseudogenes are not from repetitive regions or normal RP genes.

### Differential expression analysis

After removing the low-count genes (mean row sum ≤5 counts across all samples), we used Deseq2 in R to assess the differential expression of the remaining genes (Table. S1) in comparisons between the control and human MDD groups and between the control and chronic stress exposed mouse, divided by sex. For the GSE102556 and GSE80655 datasets, we used the SVA package in R to identify two surrogate variables that accounted for an unknown source of variation in all MDD-related contrasts (i.e., the comparison between MDD and control states in both sexes). We identified these variables by creating a full model matrix that included adjustment variables and variables representing the phenotype (i.e., control and MDD groups) as well as a null model that contained only adjustment variables. Depending on the dataset, the adjustment variables included age, RNA integrity (RIN), postmortem interval (PMI), and Ph, which were prioritized based on the total variability they explained as determined by the variance partitioning package in R (see Fig. S3). We used a p-value threshold of 0.05 for all MDD-related contrasts and false discover rate (FDR)-corrected p-value (q-value) threshold of 0.05 for all CVS-related contrasts.

### Enrichment of gene families in differentially expressed genes

To identify the stress-related ortholog genes shared by the two species (CVS: mouse and MDD: human), we performed hypergeometric overlap-based enrichment analysis implemented by the GeneOverlap package in R-3.6.0. The significance of the overlap between two gene lists can be tested using a hypergeometric distribution performed with a genomic background representing the universe of known genes (21,196 genes, default used by the package). The GeneOverlap class formulates the problem as testing whether the two gene sets are independent and then uses Fisher’s exact test to find the statistical significance. The significant overlap (q-value < 0.05) of the differentially expressed genes identified in the MDD and CVS contrasts was tested against gene family-specific gene sets (Table S2) common to both species available from the Human Genome Gene Nomenclature Committee (5). To compare the effect of gene families across the different MDD and CVS contrasts, the -log10(q-value) was used.

### Pathway enrichment analysis

To identify the biological pathways affected by the downregulated RPGs and upregulated RP-pseudogenes, we searched for the enrichment of different Gene Ontology (GO) terms associated with the identified RPGs and RP-pseudogenes in the Biological Pathway (GOBP), Molecular Function (GOMF), and Cellular Component (GOCC) categories using the GO database. RPG-pseudogene symbols for both species resemble their respective parent RPG symbols, except for an additional suffix “-ps” for mouse and “AS” for human, which we removed to perform RP-pseudogene specific enrichment analysis. Only FDR-corrected pathways (q value < 0.05) were considered for subsequent analysis. To reduce and catalogue the long list of GO terms into an interpretable format, we clustered the pathways based on biological themes as described in our previous works (10, 14, 48, 49). Briefly, the theme for a given pathway was selected based on either a text search, in which the name of the theme was used as a keyword for text query, or on the parent–child association between the GO terms in our list of significant pathways (child pathways) and the handpicked pathways in the GO database (parent pathway) representing the theme using GOdb in R.

### Seeded gene network analysis

The downregulated RPGs shared by the mouse and human datasets were considered “seed RPGs” and used to find spearman-correlation with the expression profiles of individual genes in the normalized CVS and MDD datasets (Table S3). Genes both negatively and positively correlated with a seed RPG were identified using a p-value threshold of < 0.05 and used to perform pathway enrichment analysis as described above. Both the r- and p-values were computed using the cor.test() function in the stats R package.

### Primary culture and neuron treatment

Dissociated embryonic day 18 (E18) mouse cortical cells (C57EDCX, BrainBits) were obtained from BrainBits (Springfield, IL) in 2 mL NBactive1 medium (2% B27 and 0.5 mM Glutamax). We transferred the cell medium (1 mL) into a 10-cm plate, triturating (5 x) it through a sterile P1000 micropipette tip. We dissociated the cells in the remaining 1 mL medium via trituration (5 x) in the vial. We combined the triturated media (2 mL) in a 15-mL conical tube, followed by centrifugation for 1 min at 1100 rpm (220 x g). We aspirated the supernatant with a pipet (5 mL), resuspended the cell pellet in NBactive1 medium (NB1, BrainBits), and triturated and counted the cells. We diluted cells to 1.75 ×10^5^ /mL following the manufacture’s protocol and plated 2 mL of cell suspension on a 6-well (3.5×10^5^) poly-D-lysine-coated plate (354413, Corning). Cells were incubated in a humidified environment at 37°C, 5% CO_2_ for 10 days with half of the growth medium replaced every four days. We prepared stock solutions of 1 mg/mL of dexamethasone (DEX, D4902, Sigma Aldrich) a glucocorticoid receptor agonist and 1 mg/mL mifepristone (RU-486, M8046, Sigma Aldrich), a glucocorticoid receptor antagonist, following the manufacture’s protocol. On the eleventh day *in vitro* (DIV), the growth medium was aspirated and replaced with DEX medium (2 mL, 1 μM final concentration of DEX) (18, 50), DEX and RU-486 medium (2 mL, 1 μM final concentration of DEX and RU-486 respectively), or fresh growth medium and incubated in the same environment for 72 h. The treatments were replenished at 48 h post treatment.

### RNA extraction and sequencing

Using the RNeasy Mini Kit (Cat. 74104, Qiagen), we isolated total RNA from 12 experimental replicates of the DEX, DEX+RU-486, and control groups, for a total of thirty-six samples, according to the manufacturer’s instructions. DNase I treatment (Cat.79254, Qiagen) was performed during the RNA isolation procedure to remove genomic DNA, following the manufacturer’s recommendation. Total RNA was quantified using the NanoDrop One (Thermo Scientific, USA) and Agilent Tape Station. We prepared a total RNA library for each sample, performed paired-end next-generation RNA sequencing with a depth of 60 million reads per sample on the NovaSeq platform (Illumina) at the University of Michigan Advanced Genomics Core, and processed the fast.qz files using the pipeline described above.

### Immunofluorescence analysis

To evaluate stress granule (SG) formation, we quantified the expression of the SG protein, G3BP1. We cultured mouse cortical neurons on poly D-lysine-coated glass coverslips (Neuvitro, WA) for 10 days, replacing half of the growth medium every four days. We treated the neurons with DEX or DEX + RU-486 for 72 h, fixed them with 4% paraformaldehyde (Boston Bioproducts, MA) for 15 minutes, and permeabilized them with 0.2% Triton X-100 in PBS for 15 minutes. Cells were then blocked with 3% BSA and 0.02% Tween 20 for 1 h at room temperature and incubated overnight at 4°C with anti-G3BP1 antibodies (1:250; SC-81940, Santa Cruz Biotechnology, CA), followed by a 1-h incubation with an Alexa488-conjugated anti-immunoglobulin secondary antibody (Molecular Probes) at room temperature. Cell nuclei were stained with Fluoro-Gel II with DAPI (EM Sciences, PA). We performed fluorescence and confocal microscopy assessments using a Leica CS SP5 multi-photon laser scanning confocal microscope (Leica Microsystems) and all subsequent image analysis and processing using the Leica application suite AF software (Leica Microsystems). Cells containing stress granules (SGs; n>5) >0.6μm in diameter were considered for analysis. The percentage of SG-containing cells were calculated in at least five random fields from a minimum of 30 cells per treatment. Subsequent image analysis and processing were performed using Image J software (NIH, MD). We assessed the statistical difference in stress granule formation between control, DEX, and DEX + RU-486 using one-way analysis of variance, with a p-value of <0.05 considered statistically significant.

### Quantitative PCR

We performed a qPCR assay on primary cell culture samples, which included control cells, cells treated with DEX, and cells treated with DEX + RU-486, and human postmortem ACC samples which included control and MDD subjects. Gene-specific primers for both mouse and human genes, along with the endogenous control gene *Actb*, were ordered from Integrated DNA Technologies, Inc. (IDT, IA), and the primer sequences for all analyzed genes are provided in Table S4. For the assay on primary cell culture, we selected a total of 5 genes as markers for CVS-exposed mice based on a literature review, and for humans, fifteen RPGs common between the CVS and MDD groups were used. For primary cell-culture samples total RNA was isolated from using Trizol reagent (Invitrogen), following the manufacturer’s instructions for cells cultured on poly-D-lysine-coated glass coverslips. We performed reverse transcription and cDNA synthesis using oligo-dT primers and RevertAid Reverse Transcriptase (Thermo Fisher Scientific). Gene expression for the 5 marker genes was quantified by qPCR using SYBR Green PCR Master Mix (Bio-Rad Laboratories Inc., Hercules, CA, USA), in three independent samples per treatment. For human samples, cDNA from the human postmortem ACC of control and MDD subjects was obtained from the remains of our previously published work (14), and gene expression for fifteen common RPGs was quantified using SYBR Green PCR Master Mix in six samples each for the control and MDD conditions. The statistical difference between control, DEX, and DEX + RU-486 treated primary cells was assessed using a one-way analysis of variance, and the difference between human control and MDD was assessed using a one-tailed t-test. A p-value of <0.05 was considered statistically significant.

## Data availability

The generated datasets of this study were made publicly available at the GEO repository (GEO accession number: GSE229905) and can be accessed via the following link: https://www.ncbi.nlm.nih.gov/geo/query/acc.cgi?acc=GSE229905. Codes for all bioinformatics-based analyses are provided in the associated GitHub page: https://github.com/MEladawi/Ribosome_paper

## Acknowledgments

Work is supported by DeArce-Koch memorial grant and Interdisciplinary Research Initiation Award to RS.

## Notes

### Competing Interest Statement

The authors have declared no competing interest.

