## Supplementary Material for "Ribosomal dysregulation: A conserved pathophysiological mechanism in human depression and mouse chronic stress"

**Supplementary table-legends and notes:**

**Table S1: Differentially expressed genes (DEGs) identified in various studies investigating major depressive disorder (MDD) and experimental systems of chronic stress.** The DEGs from different studies are shown in separate Excel sheets. The human postmortem MDD data was obtained from Labonte *et al.*, Ramaker *et al.*, and Shukla *et al*. (1-3). The data for mice exposed to chronic variable stress (CVS) was obtained from Labonte *et al. (1)*, and the data for mice exposed to multimodal chronic stress (mCRS) was obtained from Weger *et al.* (4). Additionally, the table includes DEGs from mouse primary neuronal cultures exposed to dexamethasone in the absence or presence of RU-486 performed in the present study. The identified genes are considered differentially expressed based on their statistical significance (p-value or q-value < 0.05) compared to control conditions.

**Table S2: Enrichment of gene families in different discovery and validation datasets shown in Fig. 1.** The sheet named “HGNC_Human” and “HGNC_Mouse” details the gene list associated with different gene families in humans and mice, respectively. The list was curated from the HGNC database (5). The sheet named “Human_Enrichment” and “Mouse_Enrichment” shows the results of the hypergeometric analysis test used to assess the significant overlap between the “HGNC_Human” and “HGNC_Mouse” gene lists and discovery (Labonte *et al. (1)*) and validation (Ramaker *et al.* and Weger *et al. (2, 4))* datasets shown in Fig. 1. The -log10(p-value) is presented, and a value greater than 1.3 (i.e., -log10(0.05)) is considered significantly enriched.

**Table S3: Pathway Enrichment Analysis Details**
**Sheets 1 and 2** contain details of gene lists correlated with RPG-seeds in human **(hs_SeedGene-Correlates)** and mouse **(hs_SeeGene-Correlates)**, respectively.

**Sheet 3 (named Fig. 2)** provides details of the pathways shown in Fig. 2. Genes correlated with RPG-seeds were used to perform the pathway analysis. The -log10(q-value) is presented, and a value greater than 1.3 (i.e., -log10(0.05)) is considered significantly enriched. Green and yellow colors indicate pathways associated with negative and positive RPG-seed correlates, respectively.

**Sheet 4 (named Fig. 3)** provides details of the pathway shown in Fig. 3. The -log10(q-value) is presented, and a value greater than 1.3 (i.e., -log10(0.05)) is considered significantly enriched.

**Sheet 5 (named Fig. S1)** provides the results of the Gene Set Enrichment Analysis (GSEA (6)) performed with the Shukla *et al.* (3) dataset. The values in columns C and D represent the GSEA-based normalized enrichment score, which were used as coordinates to plot Fig. S1B. The truth table represents different pathway themes, such as presynapse, post synapse, adaptive and innate immunity, ribosomes, and glucocorticoid stimulus response, which are shown as colored dots and were enriched in either the episode or remission state.

**Table S4:** List of primers for qPCR quantification of ribosomal protein genes in MDD patients and CVS marker genes in mouse neuronal cell culture exposed to dexamethasone in the presence or absence of RU-486.

**Supplementary figures, legends, and notes:**

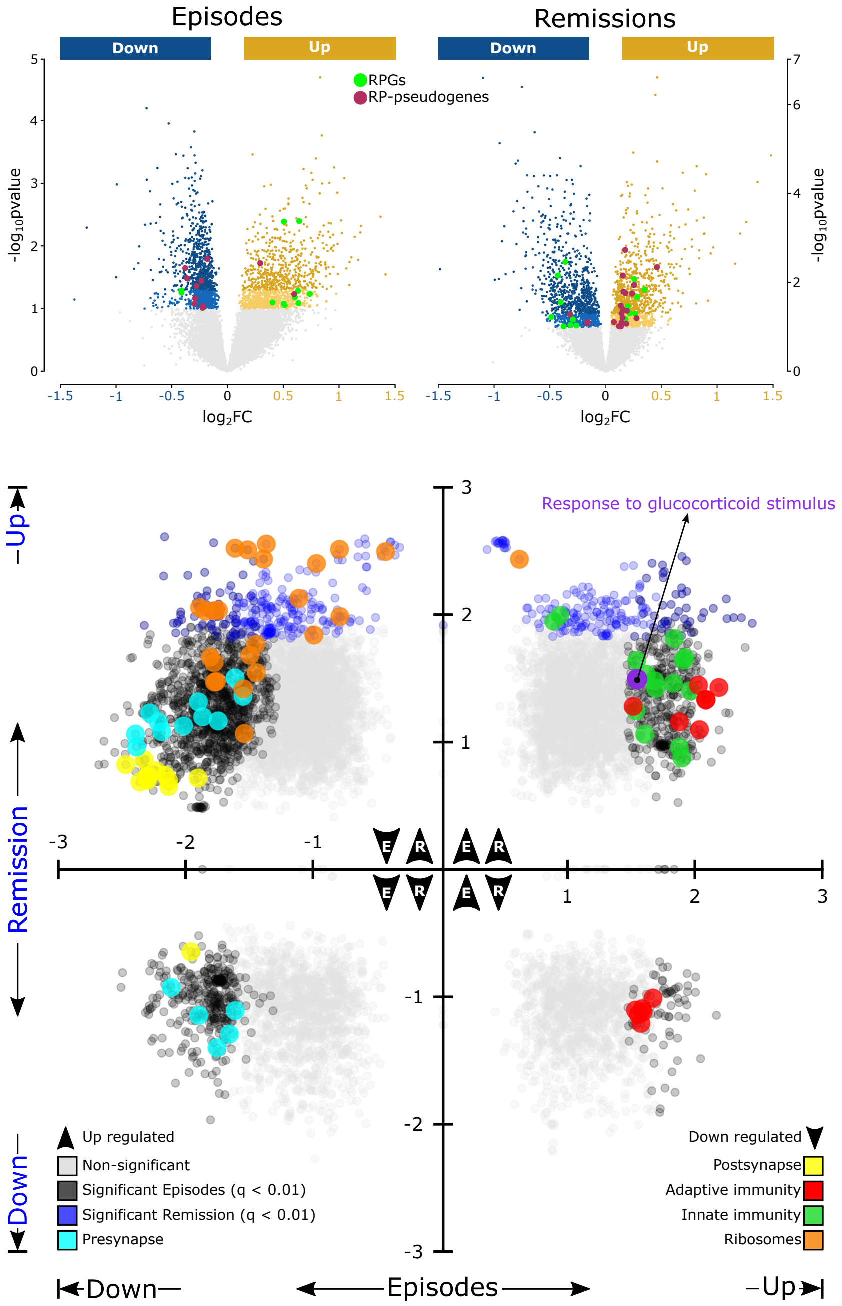

**Fig. S1: Reversibility of RPG dysregulation: A) Left:** The volcano plot shows RPG downregulation and RP-pseudogene upregulation during MDD episodes. **Right:** During remission from MDD, the pattern of RPG downregulation and RP pseudogene upregulation is reversed. As GSEA analysis (B) uses a rank-based profile to perform enrichment, it can reveal subtle changes associated with the phenotype that may not be significant at a p-value threshold of <0.05. Therefore, results with a significance threshold of >0.05 but <0.1 are also shown. **B)** Gene-set enrichment analysis of the episode (black dots) and remission state (blue dots) is shown in quadrant form. The four quadrants display distinct combinations of up- and downregulation during the two states. It is noteworthy that known synaptic and immune changes are downregulated and upregulated during episodes, respectively. Ribosome-related pathways were downregulated during the episode and upregulated during remission.

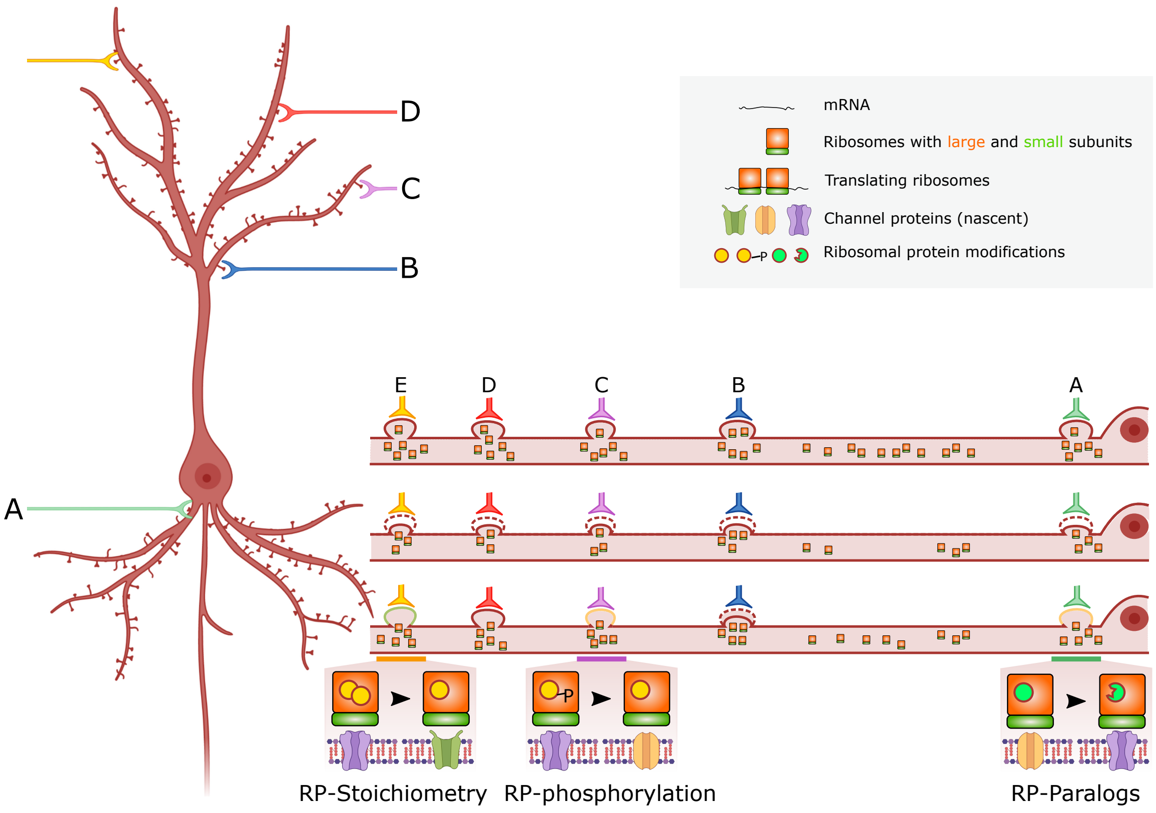

**Fig. S2:** Potential mechanisms by which RPG downregulation could affect synaptic inputs (A through E). **Global (location non-specific):** RPG downregulation may result in decreased ribosome production, which may reduce synthesis of synaptic proteins, resulting in decreased synaptic weight overall. **Local (location specific):** RPG downregulation may also change ribosome composition so that a few RPs are either removed, altered, or replaced by other RPs. These changes may result in the production of specialized ribosomes, which can alter synaptic protein translation in a cellular compartment-specific manner. Reduced ribosome production can accompany the production of specialized ribosomes, in which case reduced synaptic weight can also be observed locally.

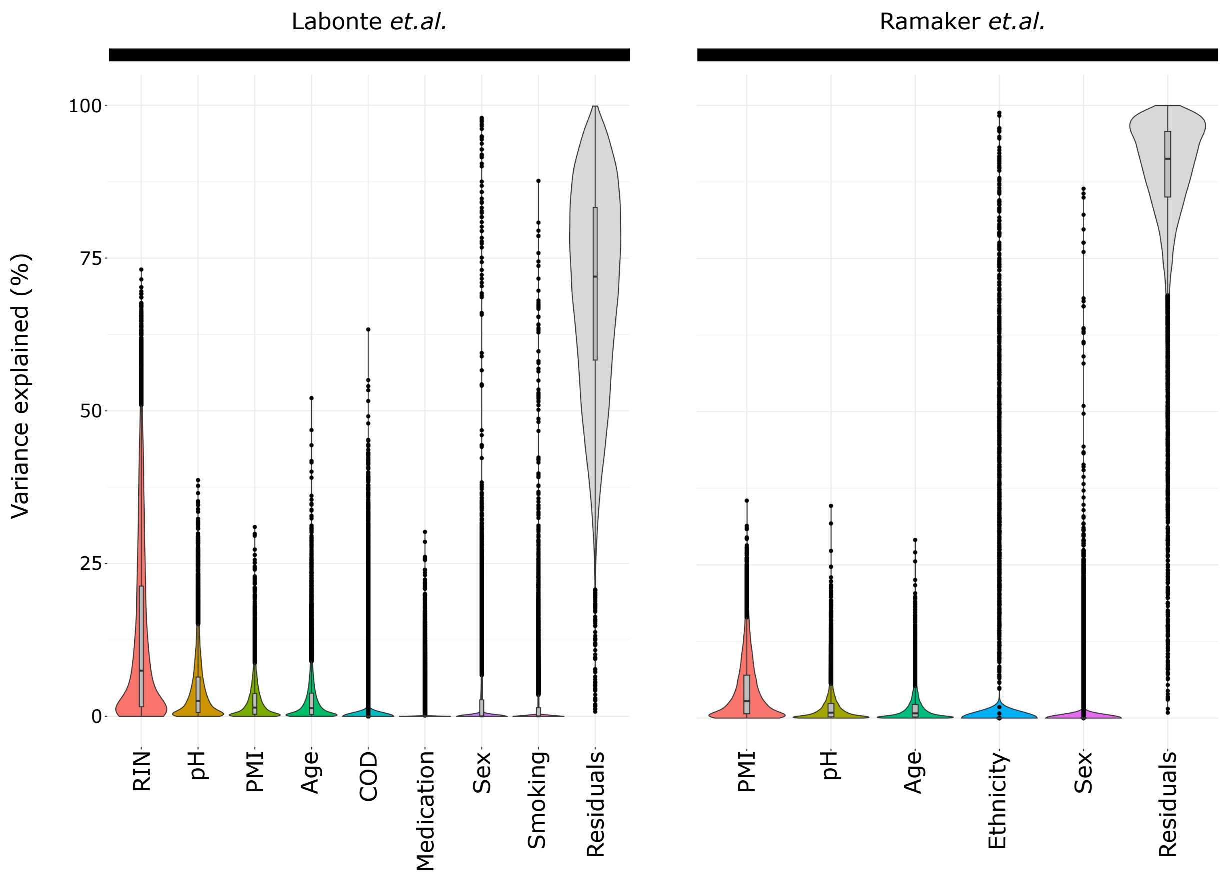

**Fig. S3: Variance explained by each variable in a gene-expression experiment.** The figure shows a genome-wide violin plot depicting the distribution of variance explained by each variable across all genes in a previously deposited dataset by Labonte *et al (1).* and *Ramaker et al (2),* using variancePartion (7) package in R. The top four variables that explain the most variation in the gene-expression profiles were regressed out during the differential expression analysis. To account for variables such as batch and medication which were not provided in the deposited data, surrogate variable analysis (SVA) package in R (8) was used to summarize these variables along with two surrogate variables that account for unknown sources of variability. RIN: RNA integrity number, PMI: postmortem interval, COD: Cause of death.
